# Gas uptake stoichiometry governs carbon partitioning in syngas-fermenting *Clostridium autoethanogenum*

**DOI:** 10.64898/2026.08.12.744430

**Authors:** Clara Vida G. C. Carneiro, Tobias Eichinger, Simra Sharif, Pratik R. Pawar, Kaspar Valgepea

## Abstract

Given the current global environmental challenges, waste biomass is an attractive renewable resource for circular economies. Gasification of biomass yields syngas (CO, CO_2_, and H_2_) that is a suitable feedstock for gas fermentation in biomanufacturing of fuels and chemicals using acetogen microbes. While it is generally known that syngas composition influences both acetogen growth and process performance, we are lacking a consistent dataset quantifying these effects under controlled fermentation conditions. Here, we mapped the metabolic response of the model-acetogen *Clostridium autoethanogenum* to seven synthetic syngas mixtures during exponential batch growth in bioreactor fermentations. Notably, distinct gas compositions resulted in different fermentation profiles, affecting both growth and metabolite production. Maximum specific growth rates ranged within 0.05 0.13 h^-1^, with slower growth for low-CO mixtures. While acetate and ethanol production yields varied between 20–133 and 76–353 mmol per gram dry cell weight, respectively, minor production of 2,3-butanediol was detected. All syngas mixtures supported co-utilization of CO and H_2_, though gas uptake stoichiometry only moderately correlated with syngas content. Importantly, gas uptake stoichiometry strongly influenced carbon partitioning, with higher relative H_2_ uptake reducing CO_2_ loss or even realizing CO_2_ fixation together with increasing carbon flow towards metabolites. Interestingly, higher syngas H_2_ content favored ethanol and 2,3-butanediol production, while higher H_2_:CO uptake ratios increased total flux through the Wood–Ljungdahl pathway rather than selectively favoring reduced by-products. Our results are valuable for a better understanding of syngas composition effects on the acetogen biocatalyst and for process engineering towards optimizing gas fermentation performance.

**Highlights:**

- Syngas composition affects acetogen growth, gas uptake, and carbon distribution
- Higher H_2_:CO uptake ratios increase carbon flow through the Wood-Ljungdahl pathway
- Higher relative H_2_ uptake reduces CO_2_ loss and increases metabolite production

## 1. Introduction

Mitigating climate change requires humankind to reduce its dependence on fossil fuel resources. Although maximal decarbonization of energy generation is the highest priority to tackle climate change, solutions for a more sustainable and fossil-free carbon-based economy in terms of fuels and chemicals also need to be deployed [1]. The future circular economies need to not only minimize waste generation but also maximize the use of renewable and waste carbon as feedstocks in industrial fuel and chemical production [2,3]. Here, non-food biomass is considered as a promising sustainable feedstock [2,4].

Among the various biomass processing technologies, gasification enables upstream methods to use nearly all biomass carbon, including fractions difficult to utilize otherwise, such as lignin [5]. Gasification with partial oxidation of biomass at high temperatures yields syngas, a gas mixture mainly composed of carbon monoxide (CO), carbon dioxide (CO_2_), and hydrogen (H_2_). Syngas composition can vary significantly depending on the biomass, feedstock, and gasification details: for instance, wood can yield ∼23% H_2_, ∼30% CO, 9–19% CO_2_ and wheat straw 48–54% H_2_, ∼30% CO, ∼14% CO_2_ [6–8]. While syngas has been widely explored as a substrate for energy generation and fuel synthesis, it has recently become an attractive substrate also for biological valorization since acetogen gas fermentation has been recently commercialized [9–11].

Acetogens are attractive microbial biocatalysts for syngas fermentation as they can naturally utilize CO, H_2_, and CO_2_ as sole carbon and energy sources through the Wood–Ljungdahl pathway (WLP) [12] and natively produce several valuable compounds, such as acetate, ethanol, and 2,3-butanediol (2,3-BDO) [11]. Among acetogens, *Clostridium autoethanogenum* is used for commercial-scale ethanol production from CO-rich waste gas streams [10] and pilot-scale production of acetone and isopropanol [13]. While the technology using CO-rich gas is well established, syngas fermentation presents new challenges. For instance, variability in syngas composition, mostly in the key components of CO and H_2_ directly affect acetogen growth, gas uptake, and product distribution, as acetogens exhibit distinct thermodynamic and enzymatic responses to different gas compositions [14,15].

In particular, the H_2_:CO ratio plays a critical role for acetogen metabolism by influencing redox balance and carbon utilization. Notably, higher H_2_ availability is expected to provide extra reducing equivalents and thus less CO would need to be oxidized and lost as CO_2_ [10,14]. Also, more substrate carbon could be directed towards reduced end-products, such as ethanol [15,16]. Indeed, supplying H_2_ to CO-fermenting chemostat cultures of *C. autoethanogenum* reduces CO_2_ loss ∼4-fold and the acetate-to-ethanol production ratio ∼5-fold [16]. In *Acetobacterium woodii*, syngas H_2_:CO ratio of ∼5.5 enhances net CO_2_ fixation and reduces acetate production compared to a ratio of ∼1.6 [17]. In contrast, higher H_2_:CO promotes acetate formation in *Clostridium ljungdahlii* [18]. Differences in the response of various acetogens to H_2_:CO further reflect variations in their bioenergetic systems and H_2_ utilization capacities [19]. These findings indicate that the relationship between H_2_:CO and product distribution in acetogens seems to be species**-**and condition-dependent, hindering our ability to predict fermentation outcomes based on syngas composition. Importantly, we are lacking a consistent dataset quantifying the effects of different syngas compositions on acetogen growth and product distribution under controlled fermentation conditions. This dataset could guide process design towards favored products [18].

Therefore, this study aimed to quantify the effects of syngas composition on the exponential growth of the model-acetogen *C. autoethanogenum* in controlled bioreactor fermentations. Batch fermentations using seven synthetic syngas mixtures revealed that higher H_2_ content promotes ethanol and 2,3-BDO production, whereas acetate formation was linked to the H_2_:CO ratio. While gas uptake stoichiometry only moderately correlated with syngas composition, it strongly influenced carbon partitioning with higher relative H_2_ uptake reducing carbon loss as CO_2_ and increasing carbon flow towards metabolites. Our results are valuable for a better understanding of how syngas composition influences both the acetogen biocatalyst and the gas fermentation process.

## 2. Methodology

### 2.1 Bacterial strain

The *C. autoethanogenum* strain LAbrini [20] deposited in the German Collection of Microorganisms and Cell Cultures (DSM 115981) was used in this work.

### 2.2 Syngas mixtures

Seven synthetic syngas mixtures (AS Linde Gas) with different proportions of CO, CO_2_, H_2_, and Ar were used (Table 1). Argon was used as the inert gas component instead of N_2_ to allow accurate bioreactor off-gas analysis. See *Results and discussion* section 3.1 for background on the choice of syngas compositions.

**Table 1.** Characteristics of synthetic syngas mixtures used in this work.

| Syngas | Composition (%) |  |  |  | Feedstock<sup>a</sup> | Reference |
| --- | --- | --- | --- | --- | --- | --- |
|  | CO | CO<sub>2</sub> | H<sub>2</sub> | Ar<sup>b</sup> |  |  |
| #1 | 50 | 20 | 20 | 10 | -<sup>c</sup> | - |
| #2 | 36 | - | 11 | 55 | Paper-reject sludge | [21] |
| #3 | 29 | 10 | 23 | 10 | Torrefied wood | [7] |
| #4 | 21 | 18 | 27 | 34 | Wheat straw | [8] |
| #5 | 30 | 3 | 29 | 38 | Straw | [22] |
| #6 | 19 | 15 | 63 | 3 | Wheat straw | [23] |
| #7 | 7 | 8 | 38 | 47 | Plastic | [24] |

<sup>a</sup>Gasification feedstock used for syngas production in respective reference. <sup>b</sup>N<sub>2</sub> is the inert gas component in the referenced studies but was replaced here with Ar to allow accurate bioreactor off-gas analysis. <sup>c</sup>Composition not derived from experimental gasification outcome and used as an “average” syngas mixture in our previous works.

### 2.3 Bioreactor batch fermentations

#### 2.3.1 Fermentation setup and operational parameters

Bioreactor batch fermentations were carried out using a bioreactor and mass spectrometer setup described in detail previously [25]. Cells were grown at 37°C and at pH 5 in a chemically defined medium (without yeast extract) described before [26] with the exception of a doubled cysteine concentration (1 g/L cysteine-HCl·H O). Starting agitation was 200 RPM that was increased by 25 RPM every four hours until 300 RPM after which it was increased by 50 RPM until 650 RPM. Starting syngas flow rate was 25 mL/min that was increased by 10% every four hours after agitation reached 500 RPM. Fermentations were carried out with three biological replicate cultures for all syngas mixtures, except with two for syngas #1 and #6 (see Table 1). Bioreactor off-gas analysis using on-line mass spectrometry data was performed as described before [26].

#### 2.3.2 Data analysis

Production yields (mmol of product/gram of dry cell weight [gDCW]) of ethanol, acetate, and 2,3-BDO were calculated using linear regression between product concentrations (mmol/L) and biomass concentration (gDCW/L). Maximum specific growth rate (μ_max_) was calculated using linear regression (R^2^ > 0.97) between ln of culture optical density (OD at 600 nm) and sample time (h). Specific product production rates (mmol/gDCW/h) were calculated by multiplying production yields with μ_max_. Specific gas uptake and production rates (mmol/DCW/h), including the specific ethanol stripping rate, were calculated using linear regression between gas uptake or production rates (mmol/L/h) and biomass concentration (gDCW/L). All linear regressions were calculated during exponential growth using 3 5 data points indicated in Fig. 2 by white square symbols, which were selected to maximize linear regression correlations and hence data reliability. Additionally, the data range for syngas #6 was selected considering that use of data points after ∼66 h could potentially confound analysis due to the preceding short period with no acetate production. Determination of carbon recoveries and balances were carried out as described before [26]. We note that ethanol stripping and the total soluble CO_2_ fraction in culture broth were included in carbon balancing.

### 2.4 Biomass concentration and exo-metabolome analysis

Biomass concentration (gDCW/L) was estimated by measuring OD and using the correlation coefficient 0.23 between OD and biomass concentration determined previously [25] using methodology described in [27]. Exo-metabolome analysis was performed using HPLC as described before [20].

## 3. Results and discussion

### 3.1 Selection of seven synthetic syngas mixtures

We selected seven different synthetic syngas mixtures (Table 1; Fig. 1a) to quantify the metabolic response of the *C. autoethanogenum* strain LAbrini [20] to syngas composition variability using controlled bioreactor batch fermentations with continuous syngas supply. The synthetic syngas compositions were chosen based on three factors: i) to mimic syngas mixtures expected from gasification of different biomass feedstocks that are relevant locally in Estonia (e.g., wood, straw, sludge; Table 1); ii) to cover a wide range of H_2_:CO ratios, in particular to test if ratios >2 allow complete conversion of CO into ethanol as predicted by thermodynamic analysis [15]; and iii) to allow comparison of results to our previous studies of syngas growth of *C. autoethanogenum* through syngas #1 (from here, syngas mixtures are referred to in text without “syngas”). Additionally, #7 was included to test a composition with H_2_:CO>5 that can be obtained from gasification of plastic waste [24]. These seven mixtures covered H_2_:CO ratios from ∼0.3 to ∼5.4 (Fig. 1b) and a broad variability in CO (7–50), H_2_ (11–63), and CO_2_ (0–20) content (Fig. 1a).

**Fig. 1.**
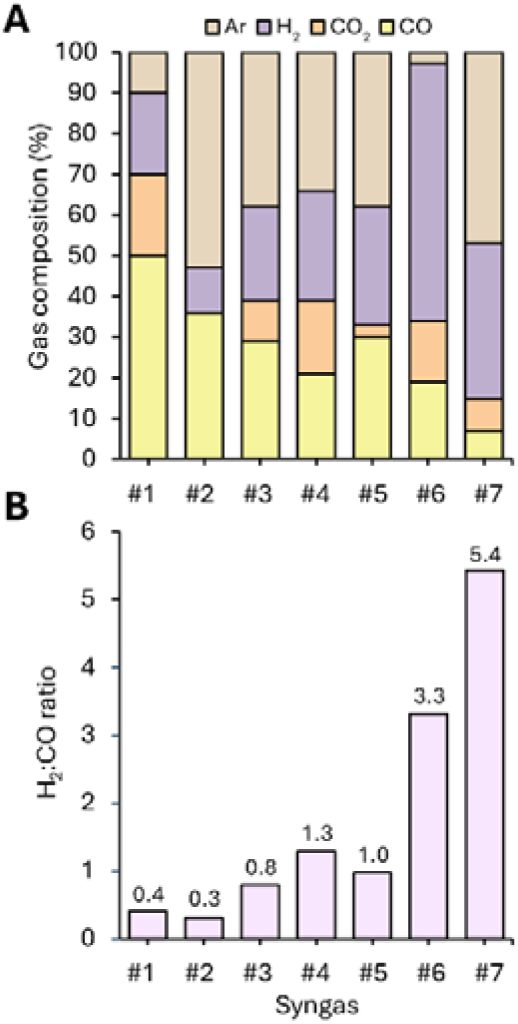
Composition of the synthetic syngas mixtures and corresponding H_2_:CO ratios used in this study. (A) Composition of syngas mixtures. Argon (Ar) was used as the inert gas component instead of N_2_ (as observed in respective gasification studies; see Table 1) to allow accurate bioreactor off-gas analysis. (B) H_2_:CO ratios for each syngas mixture.

### 3.2 Distinct effects of syngas composition on C. autoethanogenum batch growth in controlled bioreactor fermentations

We used the same gas-liquid mass-transfer profiles (agitation and gas flow) in the fermentations with all syngas mixtures to ensure data cross-comparison (see *Methodology* section 2.3.1). We note that we did not supply sulfur during the fermentation in addition to initial levels in the medium, and thus exponential growth eventually became limited by sulfur. All reported data in this study are from the period before sulfur depletion (Fig. 2a), indicated by termination of H_2_S production based on on-line off-gas analysis.

**Fig. 2.**
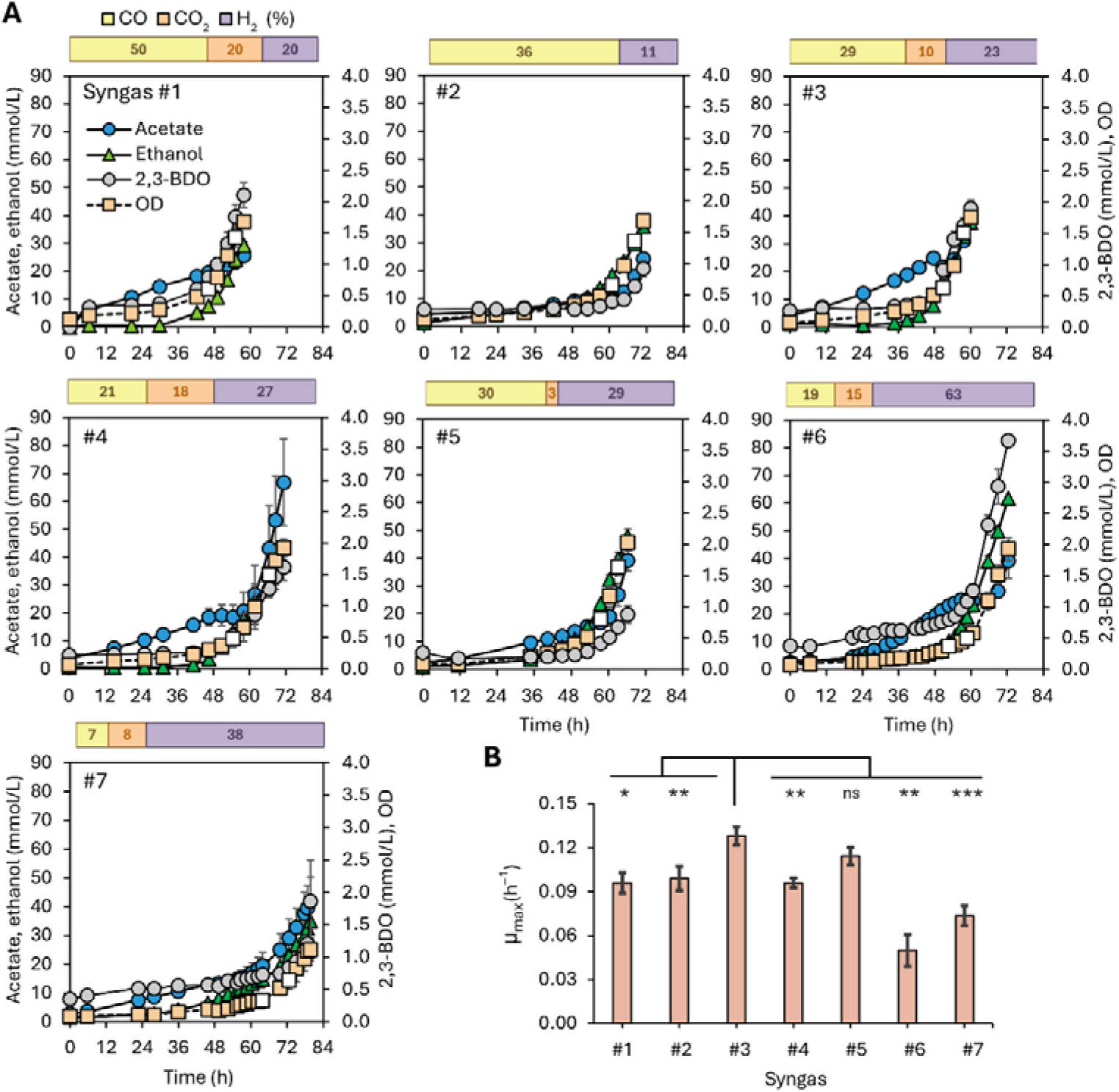
Effect of syngas composition on *C. autoethanogenum* LAbrini growth and metabolite levels in bioreactor batch fermentations. (A) Time-course profiles of growth by-product concentrations and culture optical density (OD at 600nm) for the seven synthetic syngas mixtures (% composition shown above each plot). White square symbols indicate the time range of the exponential growth phase (see *Methodology* section 2.3.2 for details); (B) Maximum specific growth rates (µ_max_) determined for exponential growth on each syngas mixture. Significance indicators: *, *p* < 0.05; **, *p* < 0.01; ***, *p* < 0.001; ns, *p* > 0.05 (two-tailed unpaired t-test). Data are presented as mean ± standard deviation between biological replicate cultures (*n* = 3, except *n* = 2 for #1 and #6). 2,3-BDO, 2,3-butanediol.

The diverse syngas compositions resulted in maximum culture ODs (OD_max_) ranging from 1.11 ± 0.17 (mean ± standard deviation) to 2.03 ± 0.09 (Fig. 2a). Assuming cultures were CO-limited, higher CO availability would be expected to support higher biomass production. While indeed the mixture (#7) with the lowest CO content (7%) achieved the lowest OD_max_, no direct positive correlation between CO% and OD_max_ was detected within the other mixtures. For instance, #2 (36% CO) and #4 (21% CO) showed similar OD_max_ and #1 with the highest CO content overall (50%) supported an OD_max_ only higher than #7 (lowest CO). Therefore, the links between syngas CO content and biomass production at least in LAbrini batch fermentations are not trivial.

We next quantified µ_max_ (Fig. 2b) for the exponential growth phase, indicated by the range between the white square symbols in Fig. 2a (*see Methodology* section 2.3.2). We note that product yields and specific rates, specific gas rates, and carbon balances are reported below also for the same exponential growth phase. LAbrini showed fastest growth on #3 (29% CO) with µ_max_ = 0.13 ± 0.01 h^-1^ being significantly higher (*p*-value < 0.05) compared to other mixtures, except for #5 (30% CO). Slowest growth was observed for mixtures with lowest CO content: #6 and #7 (Fig. 2b). This could potentially be the result of applying the same mass-transfer profile to fermentations with all gas mixtures, which could cause gas-liquid mass-transfer to limit growth at lowest CO compositions. Our results are consistent with a study using bottle batch cultures and syngas with variable CO partial pressures showing that growth of three acetogens (*C. autoethanogenum* wild-type strain JA1-1, *Clostridium carboxidivorans*, and *Acetobacterium wieringae*) is faster at higher CO levels [28]. A recent bioreactor batch fermentation study with LAbrini using a syngas composition (30% CO, 22% CO_2_, 9% H) similar to #2 in this work in terms of H_2_:CO (∼0.3) reports µ_max_ = 0.06 h^-1^ [29], which is comparable to #6 and #7 data here. These differences could possibly be explained by variable gas compositions or fermentation mass-transfer profiles.

### 3.3 Final ethanol and 2,3-butanediol levels correlate with syngas H_2_ content

While we did not operate our fermentations with the aim to achieve maximal concentrations of growth by-products, comparison of metabolite levels is still informative as syngas composition affects metabolite production [30]. In our fermentations, acetate and ethanol were the predominant products across all syngas mixtures, while 2,3-BDO was detected in much smaller amounts (Fig. 2a). Ethanol production initiated later than acetate (Fig. 2a) that is consistent with the transition from acetogenesis towards solventogenesis [31]. 2,3-BDO production increased after the exponential growth phase as seen previously for *C. autoethanogenum* [32].

A significantly higher final acetate level was measured for #4 (66.8 ± 15.4 mmol/L; *p* < 0.05), except compared to #6 and #7, while lowest levels were shown by #1 and #2 (25.6 ± 0.1 and 24.3 ± 0.9 mmol/L, respectively). The highest ethanol level was achieved by #6 (61.6 ± 0.5 mmol/L; *p* < 0.05), the syngas mixture with the highest H_2_ content (63%), and the lowest ethanol by #1 (29.4 ± 1.4 mmol/L), with similar levels for other mixtures (Fig. 2a). The highest final 2,3-BDO level was observed for #6 (3.7 ± 0.1 mmol/L; *p* < 0.05). Notably, while we measured a ∼50% higher µ_max_ of *C. autoethanogenum* LAbrini for a syngas mixture with a similar H_2_:CO ratio (∼0.3 for #2) used by Oppelt and co-workers (see above), the final ethanol and acetate concentrations compare well (∼28 and ∼22 mmol/L, respectively, for their study) [29]. *C autoethanogenum* wild-type JA1-1 batch bottle cultures using re-pressurization with a 60% CO gas mixture leads to a final acetate level of ∼27 mmol/L [28], which is similar to #1 with 50% CO in this work (∼26 mmol/L).

Interestingly, final 2,3-BDO and ethanol levels in this study showed a reasonable positive correlation with syngas H_2_ content (R^2^ = 0.67 and 0.66, respectively), but not acetate (R^2^ = 0.06) (Fig. S1). However, none of the final metabolite levels correlated with syngas H_2_:CO ratios (R^2^ = 0.07, 0.20, and 0.05 for ethanol, 2,3-BDO, and acetate, respectively). These data suggest that supply of electrons from H_2_ could affect 2,3-BDO and ethanol levels more than the balance between electron and carbon supply from syngas. This is consistent with *C. ljungdahlii* bioreactor fermentations with various syngas mixtures showing highest ethanol levels for mixtures with highest H_2_ and CO content [22]. In contrast, bottle fermentations of *C. ljungdahlii* on syngas show that higher H_2_:CO ratios promote acetate production [18].

### 3.4 Syngas H_2_ content and H_2_:CO ratio affect metabolite production yields and rates

Metabolite concentrations are dependent on biomass levels. Metabolite production yields (e.g., ethanol per biomass) account for that and are hence more reliable metrics to compare productivities across batch fermentations. Ethanol yields were higher than acetate yields for five syngas mixtures (*p* < 0.05), except for #4 and #7 (Fig. 3a; Table S1). This shows that LAbrini prefers to produce ethanol over acetate also during exponential growth, similar to steady-state growth on CO and syngas [20]. The highest ethanol yield by far (353.3 ± 17.4 mmol/gDCW) was detected for #6, which also showed the highest 2,3-BDO yield (11.9 ± 0.1 mmol/gDCW). While #6 stands out from the syngas mixtures with its highest H_2_ content (63%), the substantially higher ethanol and 2,3-BDO yields more likely result from the lowest substrate carbon flux to biomass (see Fig. 5). Higher yields for acetate were observed for #4, #6, and #7 (Fig. 3a). Interestingly, acetate yields were positively correlated with syngas H_2_:CO ratios (R^2^ = 0.71; Fig. S2). This suggests that acetate production during exponential growth is influenced not only by electron availability, but also by the balance between carbon and electron sources, likely reflecting its role as a growth-associated and ATP-generating product [33]. Specific production rates (e.g., qEtOH, mmol/gDCW/h) showed differences among syngas mixtures similar to those observed for production yields (Table S1), except for #6 displaying values closer to other mixtures (Fig. 3b) likely due to its lower µ_max_ (Fig. 2b).

**Fig. 3.**
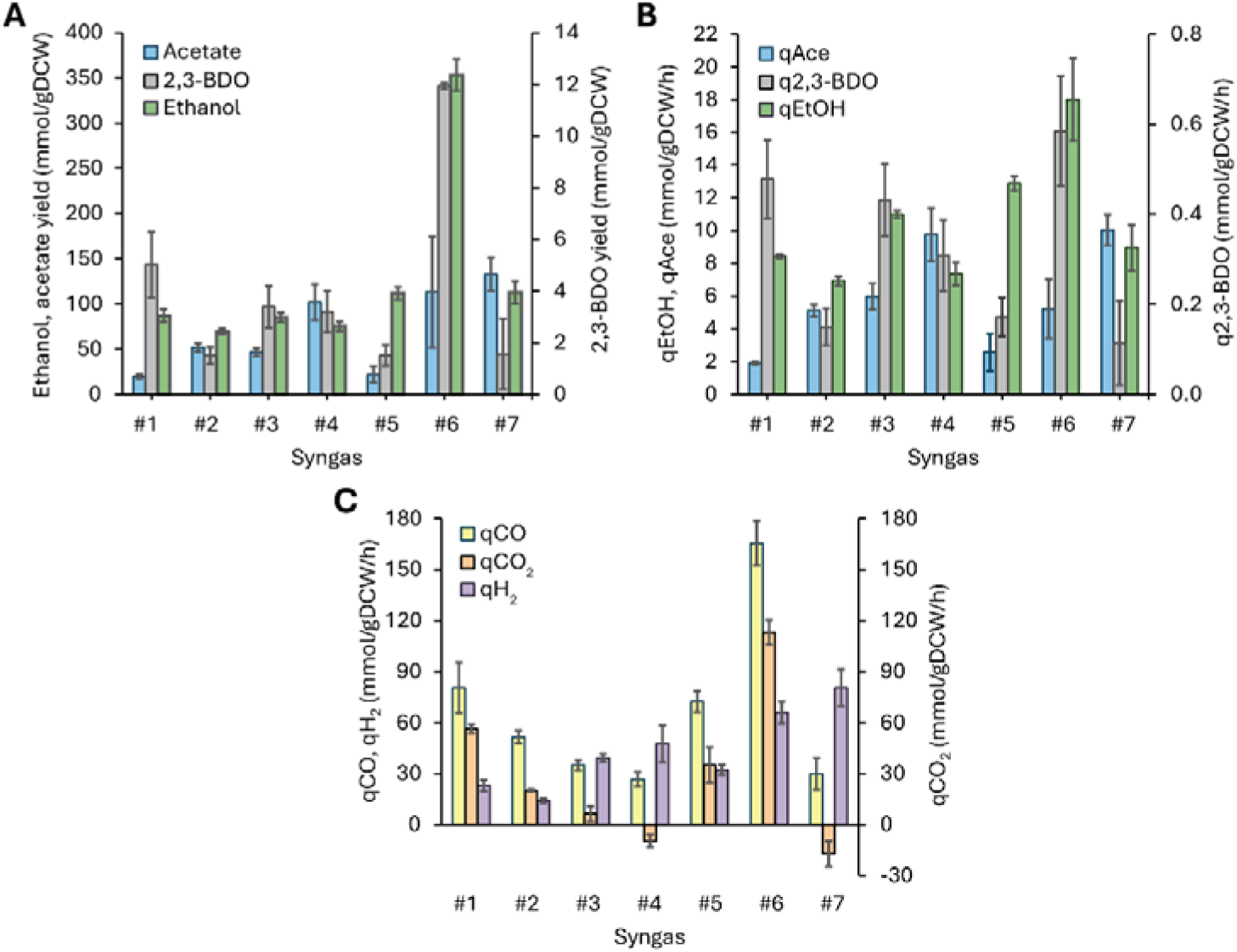
Effect of syngas composition on *C. autoethanogenum* LAbrini metabolite production and gas uptake and production. (A) Metabolite production yields. (B) Specific metabolite production rates. (C) Specific gas uptake (CO and H_2_) and production (CO_2_) rates. Negative qCO_2_ values denote uptake. Data are presented as mean ± standard deviation between biological replicate cultures (*n* = 3, except *n* = 2 for #1 and #6). gDCW, gram of dry cell weight; 2,3-BDO, 2,3-butanediol; q, specific rate.

### 3.5 Co-utilization of CO and H_2_ realizes net CO_2_ fixation in two syngas mixtures

Differences in metabolite formation suggested that carbon and electron flows varied among syngas mixtures. Thus, we next utilized our high-resolution on-line off-gas analysis to quantify gas uptake and production rates. Notably, the mixture supporting the slowest growth – #6 – showed the highest specific CO uptake rate (qCO; 165.6 ± 13.1 mmol/gDCW/h) (Fig. 3c). Similarly to its highest metabolite yields and rates (*Results and discussion* section 3.4), this can be explained by the highest H_2_ content (63%) or by the lowest substrate carbon flux to biomass in #6 (see Fig. 5). As expected, the highest specific H_2_ uptake rates (qH_2_) were measured for #6 and #7 (66.0 ± 6.4 and 80.7 ± 10.8 mmol/gDCW/h, respectively), mixtures with highest H_2_ content (63 and 38%, respectively). In general, qH_2_ showed a positive trend with syngas H_2_ content, and qCO with CO content except for #6 (Fig. S3).

Importantly, all syngas mixtures enabled simultaneous CO and H_2_ uptake by LAbrini (Fig. 3c). Co-utilization of CO and H_2_ in acetogen batch cultures is not always seen [15,16,22,34] since even low CO levels can strongly inhibit hydrogenases [35–37]. In fact, #4 and #7 enabled H_2_ uptake even higher than CO (*p* < 0.05) (Fig. 3c). Strikingly, LAbrini showed net consumption of CO_2_ for the same syngas mixtures represented by negative specific CO_2_ production rates (qCO_2_) of −9.6 ± 3.7 and −17.1 ± 7.5 mmol/gDCW/h for #4 and #7, respectively (Fig. 3c). Both theoretical stoichiometric calculations [15,34,38] and experimental data [16,33] show that acetogens need to dissipate no or less CO_2_ from CO oxidation with higher H_2_ availability. Our results further show that even CO_2_ fixation can be achieved during exponential growth on syngas when cells consume more H_2_ than CO. Intriguingly, the highest qCO_2_ of 113.1 ± 7.2 mmol/gDCW/h (i.e., CO_2_ production) was detected for the mixture with the highest H_2_ content – #6 – (Fig. 3c), suggesting that indeed very high H_2_ levels could lead to counteractive effects due to increased thermodynamic costs [15]. Within the mixtures showing CO_2_ production (positive values on Fig. 3c), qCO_2_ was higher than qEtOH and specific acetate production rate (qAce), except for #3 (Table S1).

### 3.6 Syngas H :CO ratios moderately correlate with H :CO uptake ratios

CO and H_2_ both serve as electron donors for acetogens while CO additionally provides carbon for biomass and product formation. Theoretical stoichiometric calculations with various CO and H_2_ uptake scenarios can provide estimates for carbon distribution between reduced products (e.g., ethanol) and CO_2_ for acetogen syngas fermentation [15,34,38]. However, feeding syngas to acetogens with specific H_2_:CO ratios does not guarantee proportional uptake. Quantification of the relationship between syngas H_2_:CO ratios and actual gas uptake rates is relevant not only for feedstock selection and gasification optimization, but also for the design of gas fermentation processes.

We observed a moderate positive correlation (R² = 0.50) between H_2_:CO ratios of syngas composition and uptake (Fig. 4), indicating that increased H_2_ availability is associated with proportionally higher H_2_ consumption relative to CO. The correlation is strong (R² = 0.81) if excluding #6 that deviated from the general trend (Fig. 4). The latter further suggests that additional constraints, such as increased thermodynamic costs [15] or regulatory limitations, might complicate utilization of syngas mixtures with very high H_2_ content (63% for #6). The correlation we detected could be influenced by gas-liquid mass-transfer limitations, CO inhibition of hydrogenases, CO_2_ content in syngas, or other factors. Notably, the highest H_2_:CO uptake ratio (2.9 ± 1.1) was measured for the mixture with the highest syngas H_2_:CO ratio (∼5.4 for #7).

**Fig. 4.**
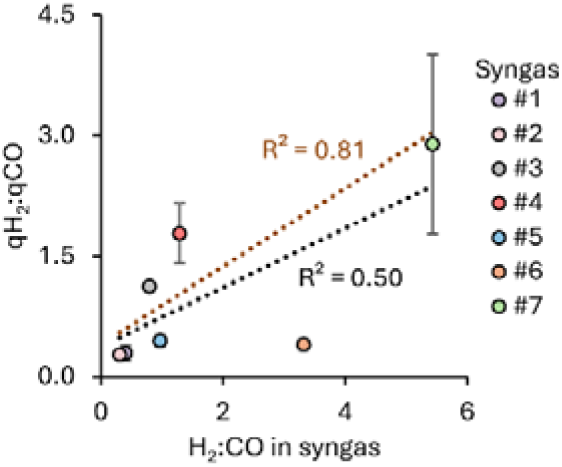
Correlation between syngas H_2_:CO ratio and *C. autoethanogenum* LAbrini H_2_:CO uptake ratio. Black dashed line is linear regression fit for all data points with corresponding correlation coefficient (R^2^) in black font; brown dashed line is linear regression fit with #6 excluded with corresponding R^2^ in brown font. Data are presented as mean ± standard deviation between biological replicate cultures (*n* = 3, except *n* = 2 for #1 and #6). q, specific uptake rate.

Our results agree with the literature showing that syngas H_2_:CO ratios influence acetogen gas uptake [14,17,18,33]. For instance, a syngas H_2_:CO ratio of 11 vs 5 increases the H_2_:CO uptake ratio from ∼8 to ∼13 in *C. autoethanogenum* steady-state chemostats [14]. Similarly in *A. woodii* steady-state chemostats, the H_2_:CO uptake ratio increases from ∼1.8 to ∼5.4 when increasing the syngas H_2_:CO ratio from ∼1.6 to ∼5.8 [17]. Interestingly, while a general linear trend is seen between the relative gas content and gas uptake also for *C. ljungdahlii* [18], CO and H_2_ uptake rates do not change between syngas H_2_:CO ratios of 1.5-to-2, potentially due to mass-transfer limitations. Taken together, syngas H_2_:CO ratios correlate with gas utilization by acetogens, while the response may deviate from linearity under conditions where gas-liquid mass-transfer or physiological constraints become limiting.

### 3.7 Syngas composition has a strong effect on carbon distribution

Quantification of substrate and product rates together with biomass formation during growth on a chemically defined medium allowed us to perform carbon balance analysis to track carbon and electron flows. After normalizing carbon recoveries to 100% to realize a fair comparison between the different syngas mixtures (see Table S2 for raw data), it was clear that syngas composition has a strong effect on carbon distribution (Fig. 5). Carbon excretion as CO_2_ was dominant in four mixtures (#1, 2, 5, 6) while none was detected in two mixtures (#4 and 7; CO_2_ was being consumed here, see Fig. 3c) and ethanol was the major carbon sink in #3. Overall, variability between the syngas mixtures was significant as carbon excretion ranged between 0 69% for CO_2_, 20 46% for ethanol, and 5 50% for acetate. Minor carbon was directed to biomass (1 11%) and 2,3-BDO (1 4%) (Fig. 5).

**Fig. 5.**
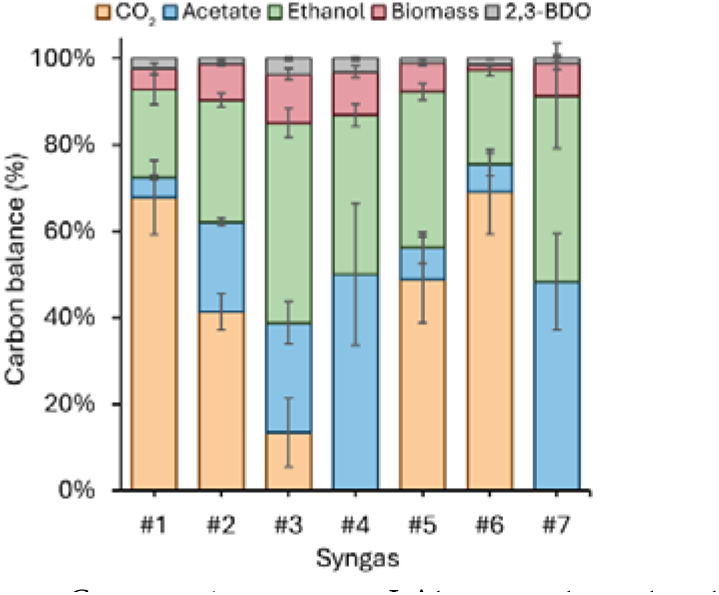
Effect of syngas composition on *C. autoethanogenum* LAbrini carbon distribution. Carbon recoveries were normalized to 100% to realize a fair comparison between syngas mixtures (see Table S2 for raw data). Data are presented as mean ± standard deviation between biological replicate cultures (*n* = 3, except *n* = 2 for #1 and #6). 2,3-BDO, 2,3-butanediol.

Notably, a substantially different carbon distribution is seen for *C. autoethanogenum* LAbrini chemostats using the same syngas as #1 in this work (50% CO, 20% CO_2_, 20% H_2_) [39]: during exponential growth in this work, most of the carbon was excreted as CO_2_ (∼68%) and ethanol (∼20%) with only ∼5% as acetate (Fig. 5), while under chemostat steady-state conditions, carbon is rather evenly distributed between CO_2_ (35%), acetate (31%), and ethanol (26%) [39]. In the *C. autoethanogenum* LAbrini bioreactor batch fermentations without yeast extract performed by Oppelt and co-workers using a syngas mixture with a similar H_2_:CO ratio (∼0.3) to #2 of this work (see above), 59% of carbon was excreted as CO_2_ [29] while the value was ∼41% for #2 (Fig. 5). These differences suggest that carbon partitioning during acetogen growth on similar syngas mixtures is influenced by both the physiological state of the culture (i.e., exponential vs gas-limited growth) and the exact gas composition and mass-transfer profiles used for batch growth.

### 3.8 Gas uptake stoichiometry governs carbon partitioning

The use of seven different syngas mixtures in this work could allow to detect potentially relevant correlations between carbon distribution and gas uptake characteristics during exponential batch growth. Indeed, our dataset revealed a positive correlation (R^2^ = 0.68) between qCO and CO_2_ % in carbon balance (Fig. 6a). Additionally, we detected a strong negative correlation (R^2^ = 0.84) between qCO and carbon flux into biomass (Fig. 6b). These observations could potentially indicate limitations in WLP throughput with increasing CO uptake. On the other hand, variable availability of H_2_ for uptake across the syngas mixtures could affect demands for CO oxidation and loss as CO_2_. Our data support the latter possibility as a negative correlation (R^2^ = 0.73) between qH_2_:qCO ratios and CO_2_ % in carbon balance was observed (Fig. 6c). This is consistent with calculations showing that higher H_2_:CO uptake ratios lead to lower excretion of CO as CO_2_ [15,34]. We previously performed *C. autoethanogenum* chemostats at different dilution rates (i.e., specific growth rates at steady-state) using syngas #1 and the same medium used here [25]. Consistent with the results here (Fig. 6a), higher qCO with faster growth in chemostats (from ∼25 to ∼73 mmol/gDCW/h) leads to an increase in carbon excretion as CO_2_ from ∼38 to ∼51%. In contrast, however, qH_2_:qCO remained unchanged at ∼0.28 between slowest and fastest chemostats, hence showing no correlation with CO_2_ production [25]. This difference further highlights the impact of the physiological state of the culture (i.e., exponential vs gas-limited growth) on acetogen phenotypes during syngas growth.

**Fig. 6.**
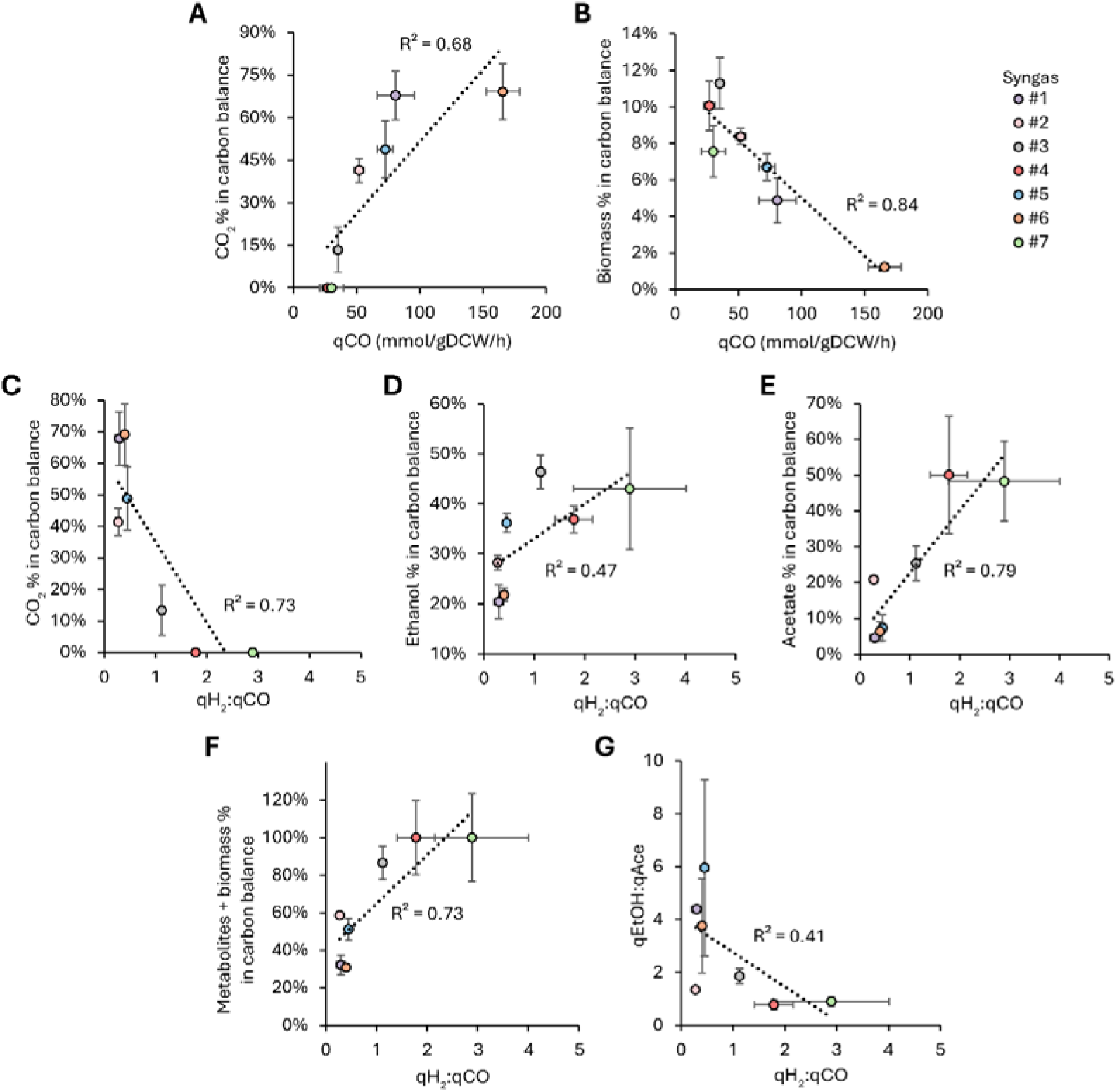
Relationship between gas uptake stoichiometries and *C. autoethanogenum* LAbrini carbon distribution. (A) Correlation between specific CO uptake rate (qCO) and CO_2_ % in carbon balance; (B) Correlation between qCO and biomass % in the carbon balance; (C) Correlation between specific H_2_:CO uptake rate ratio (qH_2_:qCO) and CO_2_ % in carbon balance; (D) Correlation between qH_2_:qCO and ethanol % in carbon balance; (D) Correlation between qH_2_:qCO and acetate % in carbon balance; (F) Correlation between qH_2_:qCO and sum of metabolites (ethanol, acetate, and 2,3-BDO) + biomass % in carbon balance; (G) Correlation between qH_2_:qCO and the ethanol-to-acetate specific production rate ratio (qEtOH:qAce). Dashed lines are linear regression fits with corresponding correlation coefficients (R^2^). Data are presented as mean ± standard deviation between biological replicate cultures (*n* = 3, except *n* = 2 for #1 and #6). gDCW, gram of dry cell weight; q, specific rate; EtOH, ethanol; Ace, acetate.

Importantly, our data show that the decreasing CO_2_ loss with increasing qH_2_:qCO leads to both higher carbon excretion as ethanol (Fig. 6d) and acetate (Fig. 6e). Furthermore, a correlation of R^2^ = 0.73 was detected between qH_2_:qCO and the total carbon allocation into products excluding CO_2_ (Fig. 6f). Thus, the extra electron supply from the higher H_2_ consumption relative to CO appears to support production of any metabolite rather than specifically favoring more reduced products. Both ethanol and acetate excretion fluxes appear not to increase further once CO_2_ excretion is avoided, around a qH_2_:qCO of 2. This indicates that the carbon split from acetyl-CoA between acetate+ethanol or biomass+2,3-BDO fluxes is rather fixed under the experimental conditions of this work (Fig. S4a).

Surprisingly, the ethanol-to-acetate production (qEtOH:qAce) did not positively correlate with higher qH_2_:qCO, a moderate negative correlation was rather observed (Fig. 6g). This is in stark contrast to *C. autoethanogenum* chemostat data showing higher ethanol-to-acetate ratios with higher H_2_:CO uptake ratios [14,16]. However, syngas H_2_:CO ratio does not affect the ethanol-to-acetate ratio in *C. ljungdahlii* batch fermentations [8]. Again, the differing phenotypic responses between exponential batch and steady-state chemostat growth potentially reflects that different regulatory principles or constraints apply on acetogen metabolism depending on the growth limitation. Therefore, cells might prioritize acetate formation during exponential growth as it is strictly coupled to ATP synthesis [40], which is expected to support biomass synthesis. Indeed, we detected a negative correlation, though rather weak (R^2^ = 0.33), between qEtOH:qAce and carbon flux to biomass (Fig. S4b).

## 4. Conclusions

Our work provides a coherent dataset quantifying the effects of syngas content on growth and product distribution of a model-acetogen under controlled fermentation conditions. We show that distinct gas compositions lead to different *C. autoethanogenum* fermentation profiles, affecting both growth and metabolite production. Our results further demonstrate that gas uptake stoichiometry governs carbon partitioning during exponential growth of *C. autoethanogenum* on syngas. More specifically, higher H_2_:CO uptake ratios increase carbon flow through the WLP rather than selectively favoring production of reduced growth by-products. Hence, metabolite production from syngas is influenced by both electron availability and the balance between electron and carbon supply. At the same time, comparison of our data with the literature highlights the impact of the physiological state of the culture (i.e., exponential batch vs gas-limited chemostat) on acetogen phenotypes. Our results are valuable for a better understanding of syngas composition effects on the acetogen biocatalyst and for process engineering towards optimizing gas fermentation performance.

## Declaration of Competing Interest

The authors declare that they have no known competing financial interests or personal relationships that could have appeared to influence the work reported in this paper.

## CRediT authorship contribution statement

Conceptualization: CV, TE, and KV; Methodology: CV, TE, SS, PP, and KV; Formal analysis: CV and KV; Investigation: CV, TE, SS, and PP; Resources: KV; Writing – Original Draft: CV and KV; Writing – Review & Editing: CV, TE, SS, PP, and KV; Supervision: KV; Project Administration: KV; Funding Acquisition: KV.

## Acknowledgements

We thank fellow group member Kurshedaktar Majibullah Shaikh for feedback on the manuscript and Karoliina Kangur and Asfand Yar Saqib for laboratory support. This work was funded by the Estonian Research Council grant RESTA9 and co-funded by the European Union and Ministry of Education and Research via project TEM-TA104.

## Notes

### Competing Interest Statement

The authors have declared no competing interest.

