## Supplementary material for "Gas uptake stoichiometry governs carbon partitioning in syngas-fermenting *Clostridium autoethanogenum*"

Carneiro *et al.*

**Table S1**

*p*-values between growth by-product metrics assessed via two-tailed unpaired *t*-test.

| **Syngas** | **Yield**  EtOH vs Ace | **q-rate**  EtOH vs Ace | **q-rate**  CO_2_ vs EtOH | **q-rate**  CO_2_ vs Ace |
| --- | --- | --- | --- | --- |
| #1 | **0.007** | **0.0003** | **0.001** | **0.001** |
| #2 | **0.006** | **0.002** | **0.00001** | **0.00001** |
| #3 | **0.0006** | **0.0004** | 0.15 | 0.85 |
| #4 | 0.09 | 0.08 | * | * |
| #5 | **0.0002** | **0.0001** | **0.02** | **0.006** |
| #6 | **0.03** | **0.03** | **0.003** | **0.002** |
| #7 | 0.19 | 0.32 | * | * |

Bold font for values denotes *p* < 0.05. Yield, production per biomass (mmol of product/gram of dry cell weight [gDCW]; q-rate, specific production rate (mmol/gDCW/h); EtOH, ethanol; Ace, acetate; ^*^net CO_2_ uptake.

**Table S2**

*C. autoethanogenum* LAbrini carbon distribution (%) in growth by-products and biomass.

| Syngas | CO_2_ | Biomass | Acetate | Ethanol | 2,3-BDO | Carbon recovery |
| --- | --- | --- | --- | --- | --- | --- |
| #1 | 70 ± 9 | 5 ± 1 | 5 ± 1 | 21 ± 4 | 2 ± 0 | 103 ± 14 |
| #2 | 39 ± 4 | 8 ± 0 | 20 ± 1 | 27 ± 1 | 1 ± 0 | 94 ± 5 |
| #3 | 18 ± 11 | 15 ± 2 | 34 ± 7 | 62 ± 4 | 5 ± 1 | 133 ± 2 |
| #4 | 0 ± 0 | 11 ± 2 | 54 ± 18 | 40 ± 3 | 3 ± 0 | 109 ± 21 |
| #5 | 48 ± 10 | 7 ± 1 | 7 ± 4 | 35 ± 2 | 1 ± 0 | 98 ± 4 |
| #6 | 69 ± 10 | 1 ± 0 | 6 ± 3 | 22 ± 1 | 1 ± 0 | 99 ± 11 |
| #7 | 0 ± 0 | 7 ± 1 | 43 ± 10 | 39 ± 11 | 1 ± 1 | 89 ± 21 |

Data are presented as mean ± standard deviation between biological replicate cultures (n = 3, except n = 2 for #1 and #6). 2,3-BDO, 2,3-butanediol.

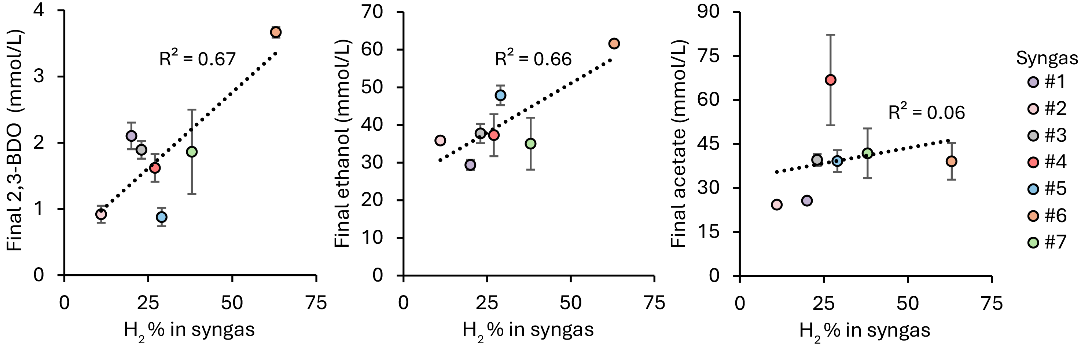

**Fig. S1.** Correlations between syngas H_2_ content and *C. autoethanogenum* LAbrini final metabolite concentrations. Dashed line is linear regression fit with corresponding correlation coefficient (R^2^). Data are presented as mean ± standard deviation between biological replicate cultures (*n* = 3, except *n* = 2 for #1 and #6). 2,3-BDO, 2,3-butanediol.

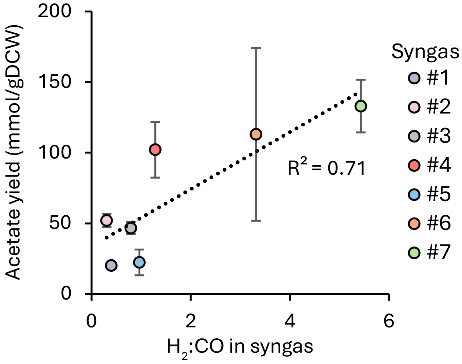

**Fig. S2.** Correlation between syngas H_2_:CO ratio and *C. autoethanogenum* LAbrini acetate production yield. Dashed line is linear regression fit with corresponding correlation coefficient (R^2^). Data are presented as mean ± standard deviation between biological replicate cultures (*n* = 3, except *n* = 2 for #1 and #6). gDCW, gram of dry cell weight.

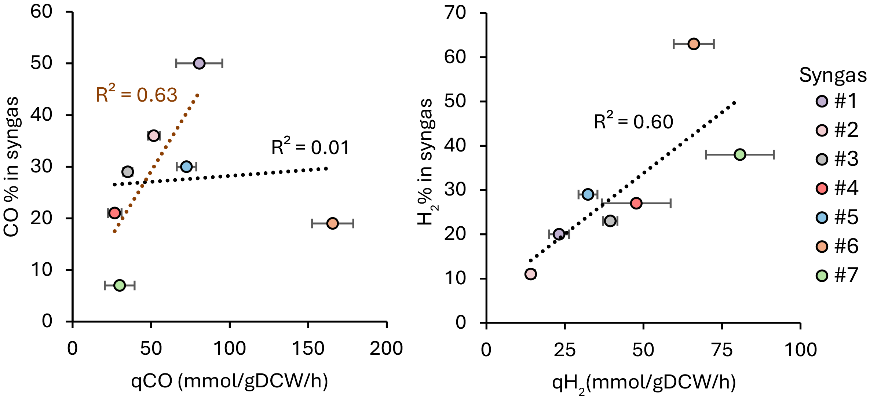

**Fig. S3.** Correlations between syngas CO or H_2_ content and respective *C. autoethanogenum* LAbrini specific uptake rate. Black dashed line is linear regression fit for all data points with corresponding correlation coefficient (R^2^) in black font; brown dashed line is linear regression fit with #6 excluded with corresponding R^2^ in brown font. Data are presented as mean ± standard deviation between biological replicate cultures (*n* = 3, except *n* = 2 for #1 and #6). q, specific uptake rate.

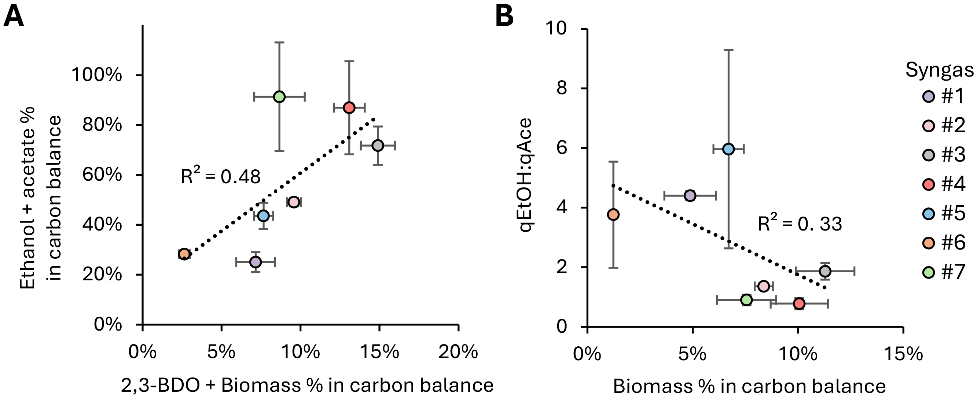

**Fig. S4.** Correlations between carbon excretion flows in *C. autoethanogenum* LAbrini. (A) Correlation between 2,3-BDO+biomass % and ethanol+acetate % in carbon balance; (B) Correlation between biomass % in carbon balance and the ethanol-to-acetate specific production rate ratio (qEtOH:qAce). Data are presented as mean ± standard deviation between biological replicate cultures (*n* = 3, except *n* = 2 for #1 and #6). EtOH, ethanol; Ace, acetate; q, specific rate.
